# Protein language model pseudolikelihoods capture features of in vivo B cell selection and evolution

**DOI:** 10.1101/2024.12.09.627494

**Authors:** Daphne van Ginneken, Anamay Samant, Karlis Daga-Krumins, Wiona Glänzer, Andreas Agrafiotis, Evgenios Kladis, Sai T. Reddy, Alexander Yermanos

## Abstract

B cell selection and evolution play crucial roles in dictating successful immune responses. Recent advancements in sequencing technologies and deep-learning strategies have paved the way for generating and exploiting an ever-growing wealth of antibody repertoire data. The self-supervised nature of protein language models (PLMs) has demonstrated the ability to learn complex representations of antibody sequences and has been leveraged for a wide range of applications including diagnostics, structural modeling, and antigen-specificity predictions. PLM-derived likelihoods have been used to improve antibody affinities in vitro, raising the question of whether PLMs can capture and predict features of B cell selection in vivo. Here, we explore how general and antibody-specific PLM-generated sequence pseudolikelihoods (SPs) relate to features of in vivo B cell selection such as expansion, isotype usage, and somatic hypermutation (SHM) at single-cell resolution. Our results demonstrate that the type of PLM and the region of the antibody input sequence significantly affect the generated SP. Contrary to previous in vitro reports, we observe a negative correlation between SPs and binding affinity, whereas repertoire features such as SHM and isotype usage were strongly correlated with SPs. By constructing evolutionary lineage trees of B cell clones from human and mouse repertoires, we observe that SHMs are routinely among the most likely mutations suggested by PLMs and that mutating residues have lower absolute likelihoods than conserved residues. Our findings highlight the potential of PLMs to predict features of antibody selection and further suggest their potential to assist in antibody discovery and engineering.

**Key points:**

– In contrast to previous in vitro work (Hie et al., 2024), we observe a negative correlation between PLM-generated SP and binding affinity. This contrast can be explained by the inherent antibody germline bias posed by PLM training data and the difference between in vivo and in vitro settings.
– Our findings also reveal a considerable correlation between SPs and repertoire features such as the V-gene family, isotype, and the amount of SHM. Moreover, labeled antigen-binding data suggested that SP is consistent with antigen-specificity and binding affinity.
– By reconstructing B cell lineage evolutionary trajectories, we detected predictable features of SHM using PLMs. We observe that SHMs are routinely among the most likely mutations suggested by PLMs and that mutating residues have lower absolute likelihoods than conserved residues.
– We demonstrate that the region of antibody sequence (CDR3 or full V(D)J) provided as input to the model, as well as the type of PLM used, influence the resulting SPs.

## Introduction

B cells recognize and bind to a variety of antigenic structures through their highly diverse and specific B cell receptors (BCRs, secreted version antibodies). BCRs are composed of two identical heavy and light chains, which consist of the variable (V), diversity (D) (only for heavy chain), joining (J), and constant (C) gene segments. The heavy chain C gene segments classify BCRs into isotypes that enable further interactions with the host immune system (1). Initial BCR diversity is generated by rearrangements of the V(D)J gene segments (2). The junction of these segments forms the third complementarity-determining region (CDR3), which exhibits the highest degree of sequence variability and contributes the most to determining antigen specificity (3). Upon antigen encounter, B cells can undergo clonal expansion and affinity maturation, where iterative rounds of SHM introduce further sequence diversity in the BCR (4). Understanding the principles of B cell selection is essential to discovering and engineering high affinity broadly neutralizing antibodies for therapeutic purposes.

Recent advancements in deep sequencing and microfluidic techniques have made it possible to comprehensively profile B cells and their corresponding antibody repertoires, providing a quantitative framework to investigate features of B cell selection (5). The rapid increase of antibody repertoire sequencing experiments has generated a wealth of data, providing fundamental insights into B cell biology and advancing applications such as antibody discovery and vaccine profiling (6–10). Furthermore, paired full-length heavy and light chain sequence information can be recovered at single-cell resolution and combined with functional characterization via recombinant expression of antibodies of interest (7). Antibody sequences can thereby be linked to biophysical properties such as antigen specificity and binding affinity. Together, such experiments have helped elucidate how single-cell repertoire sequencing can be used for antibody discovery efforts (6,9) and also shed light into fundamental insights of B cell biology pertaining to epitope convergence (9) or how transcriptional phenotypes relate to binding affinity (11).

Inspired by the recent success of natural language processing, deep learning models have been leveraged to learn complex features of protein sequences. Protein language models (PLMs) utilize unsupervised masked language modeling and are trained on large repositories of sequences. PLM representations have effectively captured structural, functional, and evolutionary features of general protein sequences (general PLM) (12–15). In line with this success, antibody-specific PLMs, trained on the massive wealth of repertoire sequencing data, have been shown to capture unique features of antibody selection such as germline gene usage and SHM (16–20). Both general and antibody-specific PLMs have been used for tasks such as humanization, structural prediction, and affinity maturation (21–23).

The masked language objective of PLMs enable the extraction of per-residue likelihoods (RLs) when supplied with unseen sequences as input. This further allows for the computation of pseudolikelihoods for entire sequences (SPs), which has been used as a computationally efficient approximation of the likelihood for entire sequences (24). It was recently demonstrated that PLM-based RLs could be leveraged to improve antibody affinity, despite remaining agnostic to affinity labels during training (21). In this work, *in silico* mutagenesis of therapeutic antibodies was performed based on PLM-suggested mutations that increased RL. In another study, the same group demonstrated that augmenting a general PLM with structural information of the entire antibody-antigen complex, using the inverse folding likelihood, can guide artificial B cell evolution (25). It however remains unknown how PLM-based likelihoods relate to in vivo features of B cell selection and evolution.

Here, we leveraged general and antibody-specific PLMs on single-cell antibody repertoire sequencing data to explore the relationship between PLM-based likelihoods and features of in vivo B cell selection and evolution. We demonstrate that the region of antibody sequence provided as input to the model, as well as the type of PLM used, influence the resulting SPs. Our findings reveal a considerable correlation between SPs and repertoire features such as V-gene family, isotype, and SHM. By reconstructing lineage evolutionary trajectories, we detected predictable features of SHM using RLs. Moreover, labeled antigen-binding data suggested that SP is consistent with antigen-specificity and binding affinity. Our study emphasizes the potential of utilizing PLMs to explore in vivo B cell selection and illustrates the power of computational tools in facilitating therapeutic antibody discovery.

## Results

### Minor correlation of SPs from varying degrees of contextual information

To study the relationship between B cell repertoire selection and SPs at single-cell resolution, we leveraged four previously published B cell repertoire datasets in the context of immunization and infection, which also provided experimental measurements of antibody specificity and affinity (Figure 1A). From these datasets, we selected samples from five mouse and five human individuals in parallel to investigate how species identity, in conjunction with the data used to pre-train protein language models, jointly influence model behavior. The mouse dataset contained BCR sequences from bone marrow plasma cells (PCs) of five mice immunized with model antigen ovalbumin (OVA) (9), hereafter referred to as Mouse1-5. The first human sample (Individual1) consisted of blood-derived B cells from an individual who had received an influenza vaccine 7 days prior to pan-B cell purification (7). The second human sample (Individual2) arises from blood-derived RBD-specific IgD-negative B cells pooled from different time points from a convalescent donor vaccinated with the SARS-CoV-2 spike protein (26). The other three human samples (Individual3-5) are part of a larger dataset that contains BCR sequences from various time points and tissues (27). From this dataset, we used three samples of blood-derived plasmablasts isolated 28 days after SARS-CoV-2 mRNA vaccination. After processing and analyzing all datasets with 10x Genomics’ Cell Ranger and the open-source R package Platypus (28), we obtained over 38,000 cells containing one heavy and one light chain. All repertoires exhibited clonal expansion for IgM, IgA, and IgG isotypes, thereby providing a comprehensive collection of repertoires to explore how PLMs can describe B cell selection (Figures 1B, S1, S2).

**Figure 1.**
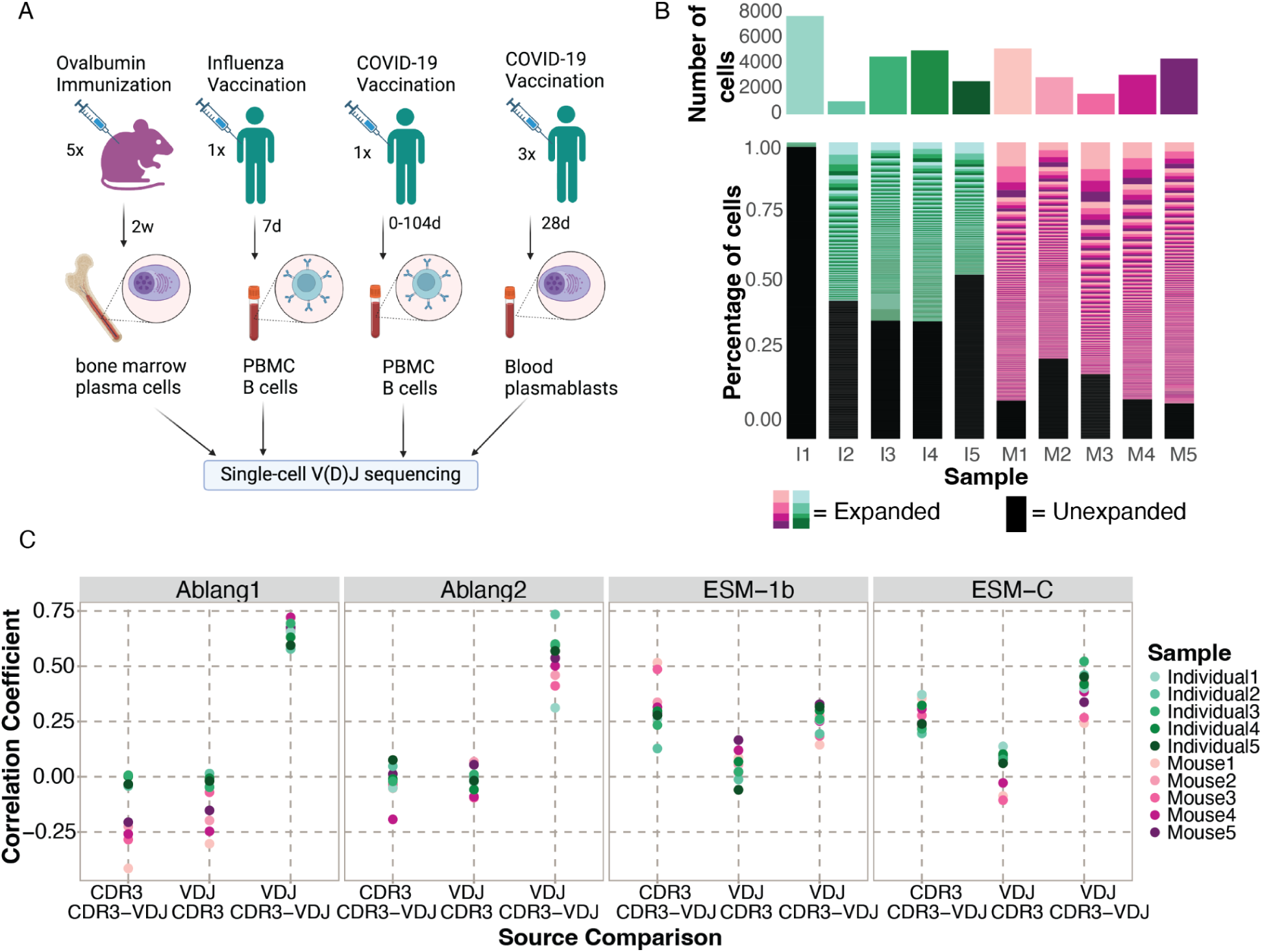
Contextual information influences SPs of antibody repertoires. A) Four publicly available single-cell V(D)J sequencing datasets with accompanying functional data were used to obtain paired heavy and light chain BCR sequences for five mice and five human samples following immune challenges. Created in BioRender. B) Total cell count (top) and distribution of clonal expansion (bottom) per sample. Each section corresponds to a unique clone, and the size corresponds to the fraction of cells relative to the total repertoire. The black color highlights the fraction of clones containing one cell. I = Individual, M = Mouse. C) Pearson’s Correlation between the SP of the heavy chains based on the input sources for four representative PLMs.

It was recently demonstrated that general PLM RLs could be employed to suggest mutations that improve antigen binding affinity by supplying full-length antibodies as input (21). We first questioned the extent to which the input sequences’ contextual information influences downstream PLM-likelihood values by calculating SPs for each antibody sequence from three input sources: 1) the full VDJ sequence for maximum contextual information (full-VDJ) 2) the CDR3 residues in the VDJ sequence for medium contextual information (CDR3-from-VDJ), and 3) only the CDR3 sequence for the minimum contextual information (CDR3-only) (Figure S3A).

We began by employing the three general PLMs ESM-C (29), ESM-1b (12) and ProtBERT (13) as well as the three antibody-specific PLMs Ablang2 (20), Ablang1 (18) and Sapiens (22) on the heavy chain sequences from all input sources for all samples. Next, we calculated the correlation between SPs from the three input sources (Figures 1C, S3B). We observed no correlation between the SPs from the CDR3-only to either CDR3-from-VDJ and full-VDJ sources for all samples (average r^2^ = 0.088 and -0.020, respectively). However, a correlation was observed when comparing the SPs from the sources full-VDJ to CDR3-from-VDJ (average r^2^ = 0.427). This correlation was especially prominent in the antibody-specific PLMs Ablang1 and Ablang2 for the human samples (average r^2^ = 0.631 and 0.562, respectively). Comparable correlations were observed when calculating the SP for the light chains (Figure S3C). Together, these results recapitulate the ability of PLMs to capture long-range sequence interactions (12,13,16,22) and highlight the importance of the full-length antibody sequence in determining SPs.

### SPs from different PLMs were marginally correlated

Different PLMs have been demonstrated to influence immune receptor representations (6,19). We therefore investigated how several general and antibody-specific PLMs correlated on in vivo antibody repertoire data for the three different input sources on the heavy chain sequences (Figure 2A, S4A). For the human samples, these SPs were slightly correlated across all models (average r^2^ =0.383), and this correlation increased significantly when more contextual information was provided as input source (CDR3-from-VDJ average r^2^ = 0.411 and full-VDJ average r^2^ = 0.614). We observed the highest correlation between the antibody-specific PLMs Ablang1 and Sapiens when using the full-VDJ input sequences of the human samples (average r^2^ = 0.860). Overall correlations were lower for the mouse samples (average r^2^ = 0.266) and did not always increase when providing more contextual information. For the mouse samples, we observed the highest correlation between the general PLMs (ProtBERT, ESM-C, and ESM-1b) when using the full-VDJ input sequence (average r^2^ = 0.688). The trend that general PLMs had the highest correlation also held true when analyzing the light chains (Figure S4B). We observed a positive correlation between heavy chain SP and light chain SP for every PLM, except for Ablang2 (Figure 2B). Ablang2, in contrast to the other five PLMs used in this work, is fine-tuned on paired heavy and light chain sequences and is consequently able to compute a paired SP of both chains. Interestingly, Ablang2 paired SP is positively correlated with Ablang2 heavy chain SP, but negatively correlated with Ablang2 light chain SP (Figure 2C). Based on these findings and the more prominent role the heavy chain plays in antigen binding and structural properties (3), we decided to focus on the SPs of the full-VDJ heavy chain sequences for the rest of the analysis for Ablang1, ESM-1b, ESM-C, ProtBERT, and Sapiens. To fully utilize the single-cell nature of our data, we use the paired SP for the rest of the analysis for Ablang2.

**Figure 2.**
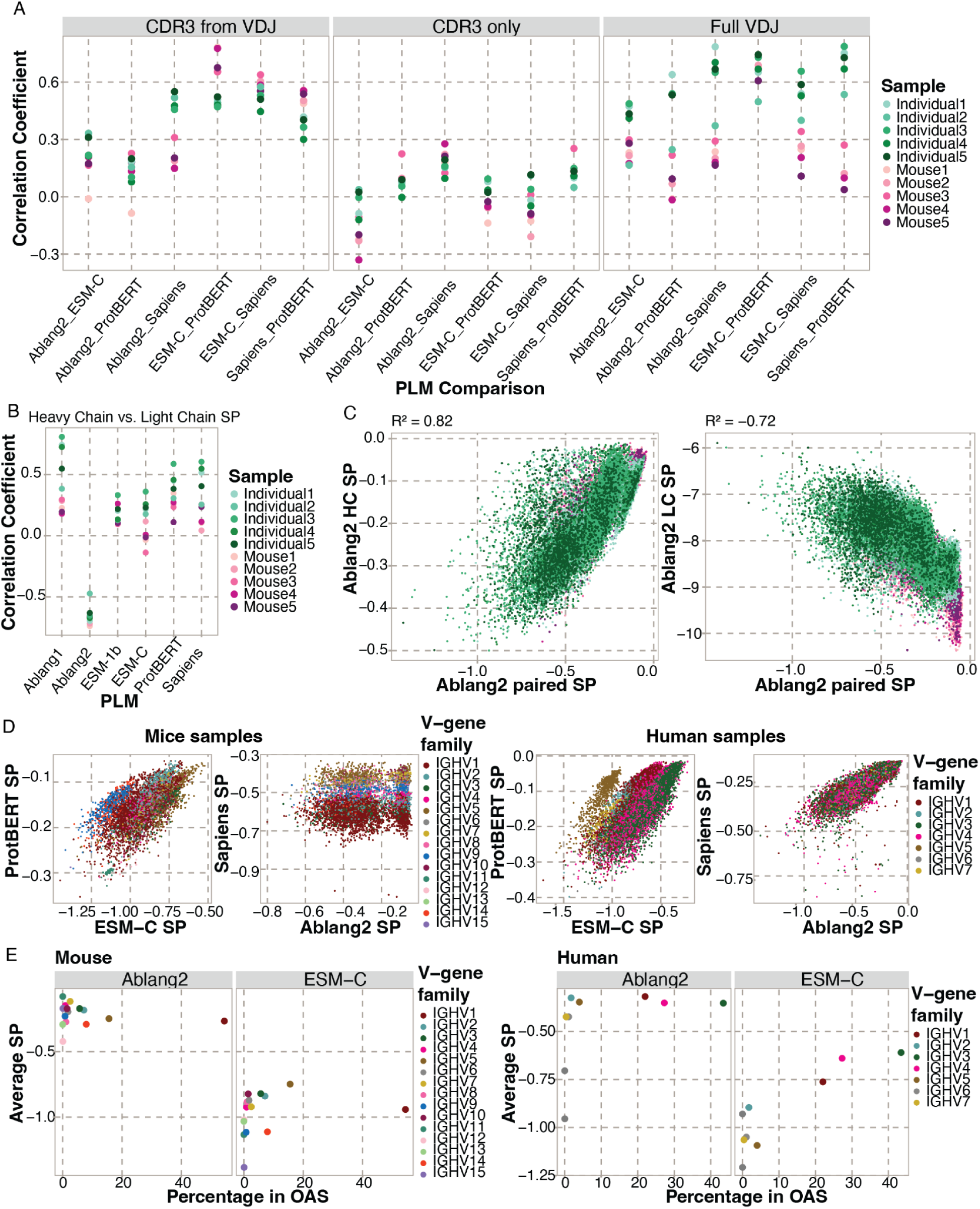
Correlation between SPs calculated with four different PLMs. A) Pearson correlation between the heavy chain SP calculated with the different PLMs for each of the three input sources. B) Correlation between the heavy and light chain SPs of the full V(D)J sequences. C) Correlation between the SPs calculated with Ablang2 based on the paired heavy and light chain (x-axis) and only the heavy chain (left y-axis) or light chain (right y-axis). D) SPs calculated with different PLMs for mice (left) and human (right) samples, colored by the V-gene family. E) The average SP per V-gene family and the percentage of unique sequences of this V-gene family in the OAS database for the mice (left) and human (right) samples.

After observing the major contribution that the full-length antibody sequence plays in computed SPs (Figure 1), we questioned whether certain V-genes were routinely deemed as more likely and whether this was consistent when using different PLMs. We therefore stratified SPs by V-gene subfamilies, which demonstrated a bias toward certain V-genes (Figure 2D). For ProtBERT, mouse sequences belonging to the IGHV1 family showed a relatively higher SP than other V-gene subfamilies (average SP of -0.159 v -0.164, respectively). This contrast was reversed for ESM-C, where mouse sequences of the IGHV1 family showed a slightly lower SP (average SP of -0.941 v -0.918, respectively). For the human samples, sequences belonging to the IGHV1 and IGHV4 families showed a lower SP for ESM-C in comparison to sequences from the other V-gene subfamilies (average SP of -0.700 v -0.675, respectively). No significant V-gene bias was observed with the other PLMs for the human samples. However, the antibody-specific PLM Sapiens showed a clear V-gene bias for the mouse sequences, with the IGHV1 family showing the lowest SP and IGHV7 the highest (average SP of -0.610 v -0.412, respectively). This observed V-gene bias with Sapiens in the mouse but not in the human samples is surprising considering Sapiens was trained solely on human antibody sequences. We next questioned whether this observed V-gene bias was correlated with the potential enrichment of certain V-genes in the training data of PLMs, such as the Observed Antibody Space (OAS) (30) for the antibody-specific PLMs. We observed no significant correlation between the OAS frequency and average SP per V-gene family for the mouse samples (average r^2^ = 0.196) (Figure 2E, S5A). In contrast, the V-gene family OAS frequencies and average SP for the human samples revealed a correlation (average r^2^ = 0.879) (Figure 2E, S5B). This correlation was more prominent for the general PLMs, despite not being trained directly on sequences from the OAS database and applying a strict sequence similarity threshold for their training data. Together, these findings demonstrate the importance of model selection when using PLM likelihoods to engineer antibodies.

### Repertoire SPs correlate with isotype usage and SHM but not clonal expansion

PLMs and associated likelihoods have recently demonstrated the ability to contextualize properties of antibodies such as germline gene usage and degree of SHM (16), but such analyses have not yet been performed to investigate selection within entire repertoires at single-cell resolution. We, therefore, questioned whether PLM-based likelihoods were correlated to certain signatures of B cell selection in the various immunization and infection settings. As the majority of training data for antibody-specific PLMs arises from the human IgM isotype (30), we hypothesized that the SPs corresponding to IgM B cells would be higher than other isotypes.

While we indeed found that the IgM isotype in human repertoires corresponds to significantly higher SP values than IgG and IgA for antibody-specific PLMs, we additionally observed that this was also true for general PLMs despite being trained on general protein corpora (average SP of –0.306 v -0.386, respectively) (Figures 3A, S6B). As Sapiens is exclusively trained on human antibodies, we questioned whether SPs of murine repertoire would demonstrate more comparable SPs between isotypes. SPs from all antibody-specific PLMs were higher for murine IgM BCRs compared to IgA and IgG (average SP of -0.387 v -0.487, respectively), however, this was not the case for every general PLM (Figures 3A, S6A). We questioned whether this observed isotype bias correlated with the potential enrichment of certain isotypes in the OAS database. As expected, the average SP per isotype was correlated with its frequency in the OAS databases (Figure 3B, 3C, S6C, S6D). Together, these findings suggest that using either general or antibody-specific PLMs for antibody engineering can bias proposed mutations towards IgM-like antibodies.

**Figure 3.**
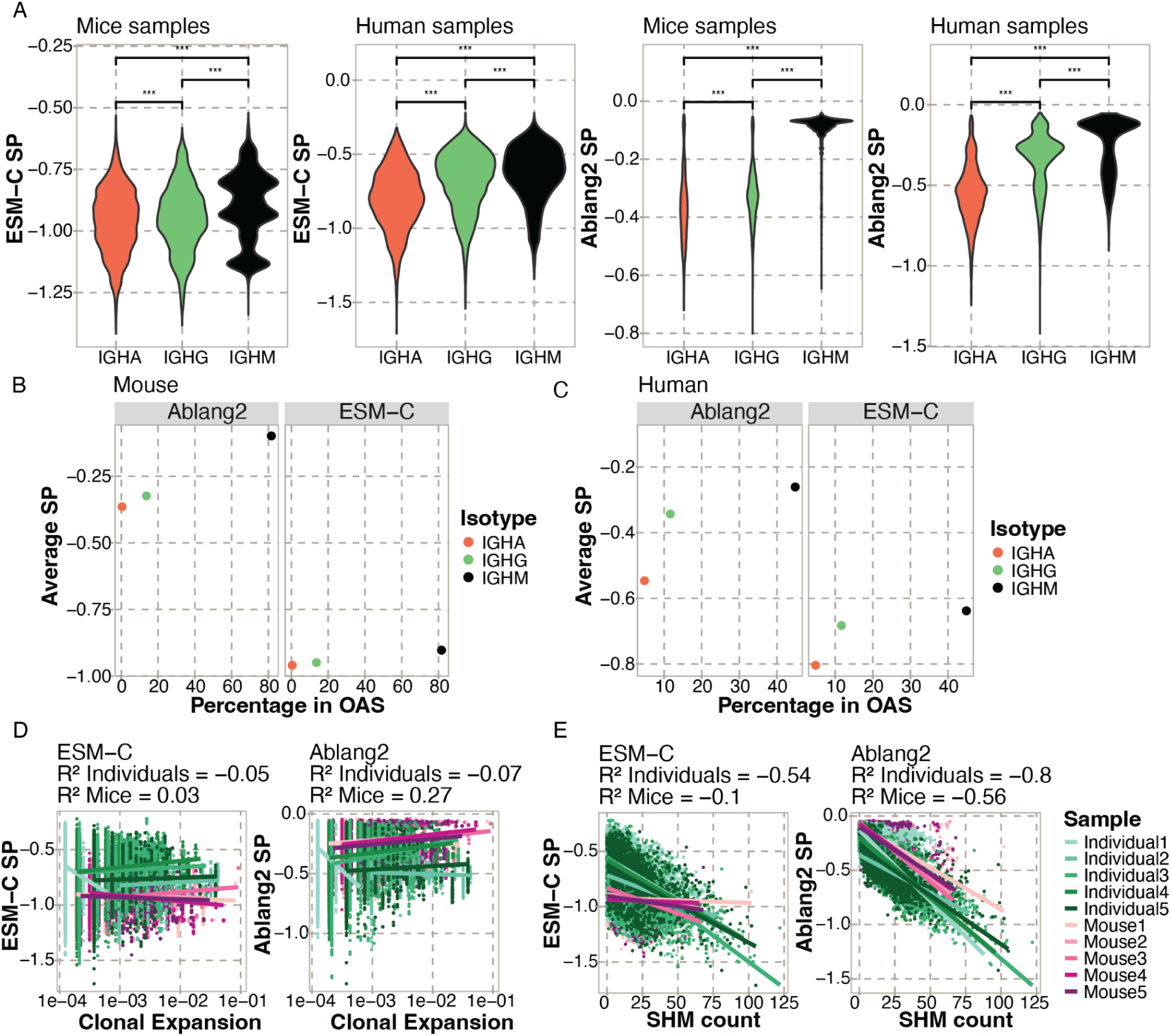
Correlation between SPs and features of B cell repertoire evolution. A) Distribution of SPs of BCRs from certain isotypes for a general PLM (left) and an antibody-specific PLM (right). T-test significance: *** = adjusted p-value below 0.001. B) The average SP per isotype and the percentage of unique sequences of this isotype in the OAS database for the mouse samples. C) The average SP per isotype and the percentage of unique sequences of this isotype in the OAS database for the human samples. D) Pearson Correlation between normalized clonal expansion (number of cells per clonotype divided by the sample size) and SP for ESM-C (left) and Ablang2 (right). E) Pearson Correlation between the amount of SHM (Hamming distance from the germline) and SP for ESM-C (left) and Ablang2 (right).

We next leveraged the single-cell nature of the repertoire datasets to question whether the degree of clonal expansion correlated with SPs. We hypothesized that antibodies deemed more evolutionarily favorable may correspond to some inherent selection bias and thereby have relatively larger degrees of clonal expansion. We computed the relative expansion within each repertoire and related this to the SPs for each PLM. We observed no major correlation between clonal expansion and SPs in humans and mice for both the general and antibody-specific PLMs (average r^2^ = -0.14) (Figures 3D, S7A). Finally, as previous studies have demonstrated that PLM-based embeddings are sensitive to the amount of SHM (16,17), we finally questioned whether the amount of SHM correlated with SPs. We observed a negative trend between SP and the extent of SHM for the human samples (average r^2^ = -0.54) (Figures 3E, S7B). This negative correlation increased remarkably when calculating SP with antibody-specific PLMs (average r^2^ = -0.8), in which case the mouse samples also showed a negative trend (average r^2^ = -0.56). Taken together, these findings reveal trends between repertoire features and SPs, suggesting a correlation between PLM-derived likelihoods and natural B cell selection.

### SHM coincides with RLs

Thus far, we have explored the relationship between SHM and SP on the level of the repertoire. However, SHM occurs within a lineage-specific context, thereby motivating us to investigate how PLM-based likelihoods can infer or predict features of SHM on the level of the individual clone. We therefore constructed lineage trees using the Levenshtein distance to represent the evolutionary trajectory of SHM within each clone (Figures 4A). For the human samples, we observed that sequences closer to the germline had an overall higher SP (average r^2^ = -0.523), this trend was less pronounced in the mice samples (average r^2^ = -0.205) (Figures 4B, S8). This observation is in line with the repertoire-wide results of SHM count. We repeated this analysis for trees constructed using a maximum parsimony algorithm and obtained comparable results (Figure S9).

**Figure 4.**
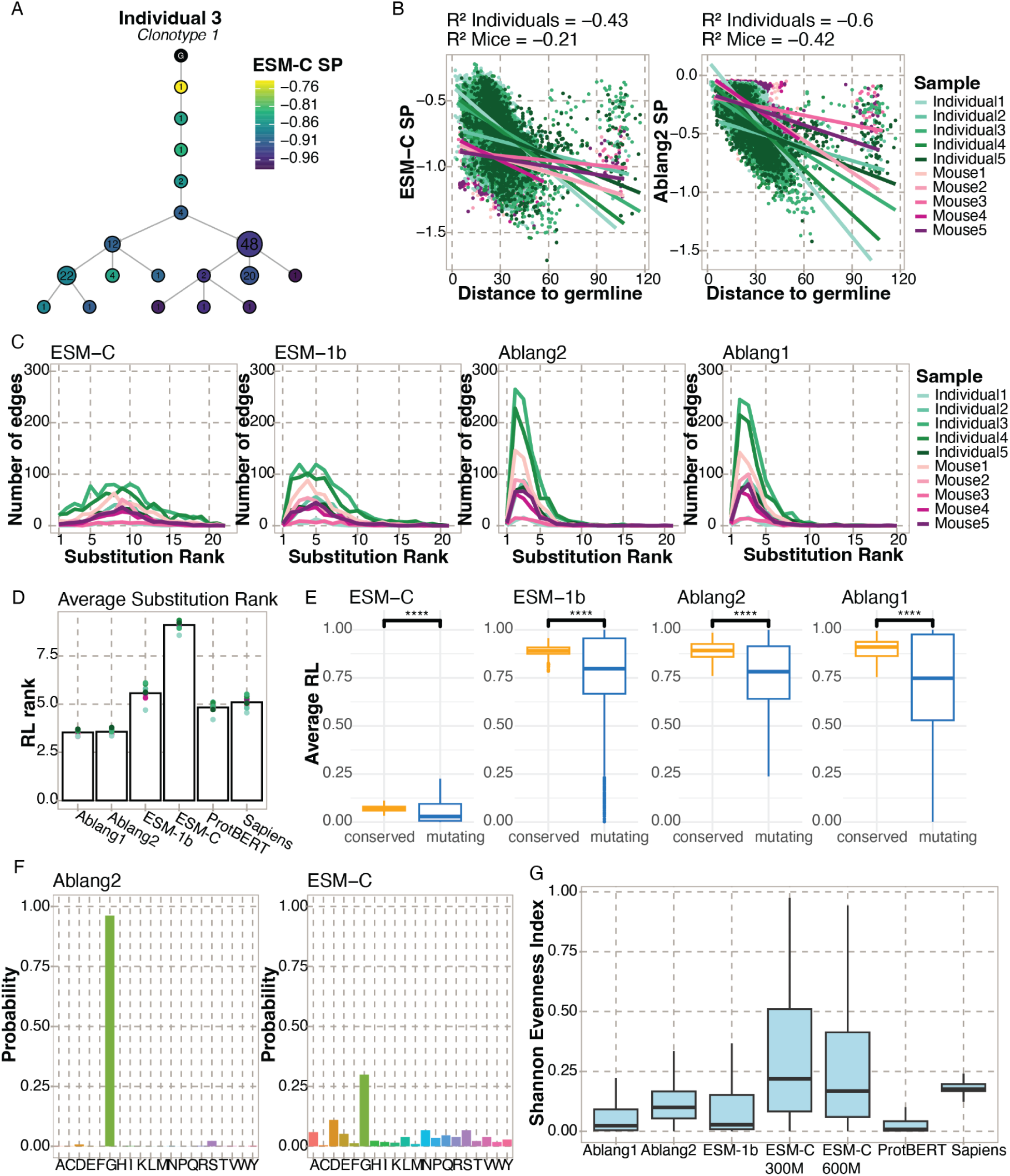
SHM coincides with pseudolikelihoods (SPs) and per-residue likelihoods (RLs). A) Representative lineage tree colored by ESM-C heavy chain SP. Numbers in the nodes indicate the number of cells. B) Pearson Correlation between ESM-C and Ablang2 SP and the total edge length to the germline (Levenshtein distance) for each sequence in all trees. C) The RL ranks of the substitutions along the edges of the lineage trees. The average rank is used for edges with multiple substitutions. D) Mean substitution RL rank for each sample (dots) and average of all samples (bars) per PLM. E) Difference in average RL between conserved and mutating residues. T-test significance: **** = adjusted p-value below 0.0001. F) Example distribution of the residue likelihoods for one position in a BCR sequence for Ablang2 and ESM-C. G) The Shannon Evenness Index of the RL distributions of all positions in all sequences per PLM.

Next, we focused on the substitutions along the edges of the trees and the corresponding RL rank of those mutating positions. Edges in the trees usually represented a mutation toward one of the most likely residues at that position, except for the most likely (Figure 4C). Ablang1 and Ablang2 substitution ranks were on average the highest, as expected given their training data explicitly includes mutated BCRs (Figure 4D). To investigate the predictability of the location of SHM, we compared the original RL of the mutating residues with the non-mutating conserved residues along the edges, which demonstrated that mutating residues corresponded to a lower RL than conserved residues (Figure 4E). For ESM-C, we observed significantly lower RLs than for the other PLMs and a higher average substitution rank. We questioned whether ESM-C fails to capture SHM, or whether the RL distribution per position diverges from those of other PLMs. We observed that ESM-C captures the same patterns as other PLMs, such as which amino acid is most likely, however, this was routinely predicted with lower likelihood values (Figure 4F). We quantified this by calculating the Shannon Evenness Index of PLM probabilities for every position in every sequence. This revealed a more even distribution for ESM-C in comparison to the other PLMs, even when increasing the number of parameters for ESM-C. We finally repeated these analyses in additional repertoires to assess the generalizability of these findings. Indeed, we observed that SHMs were ranked among the most probable by PLMs and that mutating residues had a lower RL than non-mutating residues (Figure S10). Together, these results indicate that PLM-based likelihoods negatively correspond with reconstructed B cell evolution, as both SPs and RLs decrease along the edges of the trees. From the mutating residues, we can conclude that RLs can predict, to some extent, to what amino acid SHM will mutate and at which position.

### SPs were correlated with functional features of antibodies

Having observed that SPs correlate with B cell repertoire features, we next questioned whether this relationship would also be reflected on a functional level. To this end, for the OVA-immunized mouse dataset, we considered OVA specificity labels as tested by ELISA for the most abundant sequences in the 177 most expanded IgG clones (9). A high degree of stochasticity in antigen binding was previously reported, as binding and non-binding clones were evenly distributed at all levels of expansion in the repertoire. For the human dataset of Kim *et al.* (Individual3-5 here), antigen specificity to SARS-CoV-2 S was tested by ELISA on 2,099 recombinant monoclonal antibodies derived from eight donors (27). These antibodies were generated from expanded clones in lymph node samples at seven and fifteen weeks post-vaccination. From these antibodies, 71.6% were shown to bind to the spike (S)-protein. We next questioned whether SPs could be used to predict the selection of binding clones and therefore aid in the discovery of therapeutic antibodies. The results varied depending on the PLM: while ESM-1b exhibited higher SPs for binders, other models showed no significant difference, and notably, Sapiens reported lower SPs for binders (Figure S11A). For the human antibodies, SPs calculated with all PLMs were significantly higher for binders than for non-binders, however the differences were minor (average SP of -0.317 v -0.337, respectively) (Figure S11C). These results suggest that SPs may capture features of in vivo antigen-specificity in the context of immunization, but SP alone is insufficient to discover binding antibodies.

It was previously demonstrated that antibody affinity was related to RLs in in vitro engineering efforts (21). We therefore questioned whether this relationship was also present in the context of in vivo B cell selection. For 58 of the OVA-specific antibodies from the mouse dataset, the affinity to their cognate antigen was previously measured using biolayer interferometry (BLI) (9). No correlation was found between binding affinity (determined by the dissociation constant (Kd)), clonal expansion, and the amount of SHM. For the human dataset of Kim *et al.* (27), binding affinity to the S-protein was measured with BLI for 43 antibodies derived from clonally related plasmablasts and bone marrow PCs from seven individuals. In our analysis, we observed no strong correlation between binding affinity and SPs for all PLMs. For the mice dataset, the highest correlation was observed for ESM-C (r^2^ = 0.212) (Figure 5A, Figure S11B). For the human data, the highest correlation was observed for ESM-1b (r^2^ = 0.378) (Figure 5B, Figure S11D).

**Figure 5.**
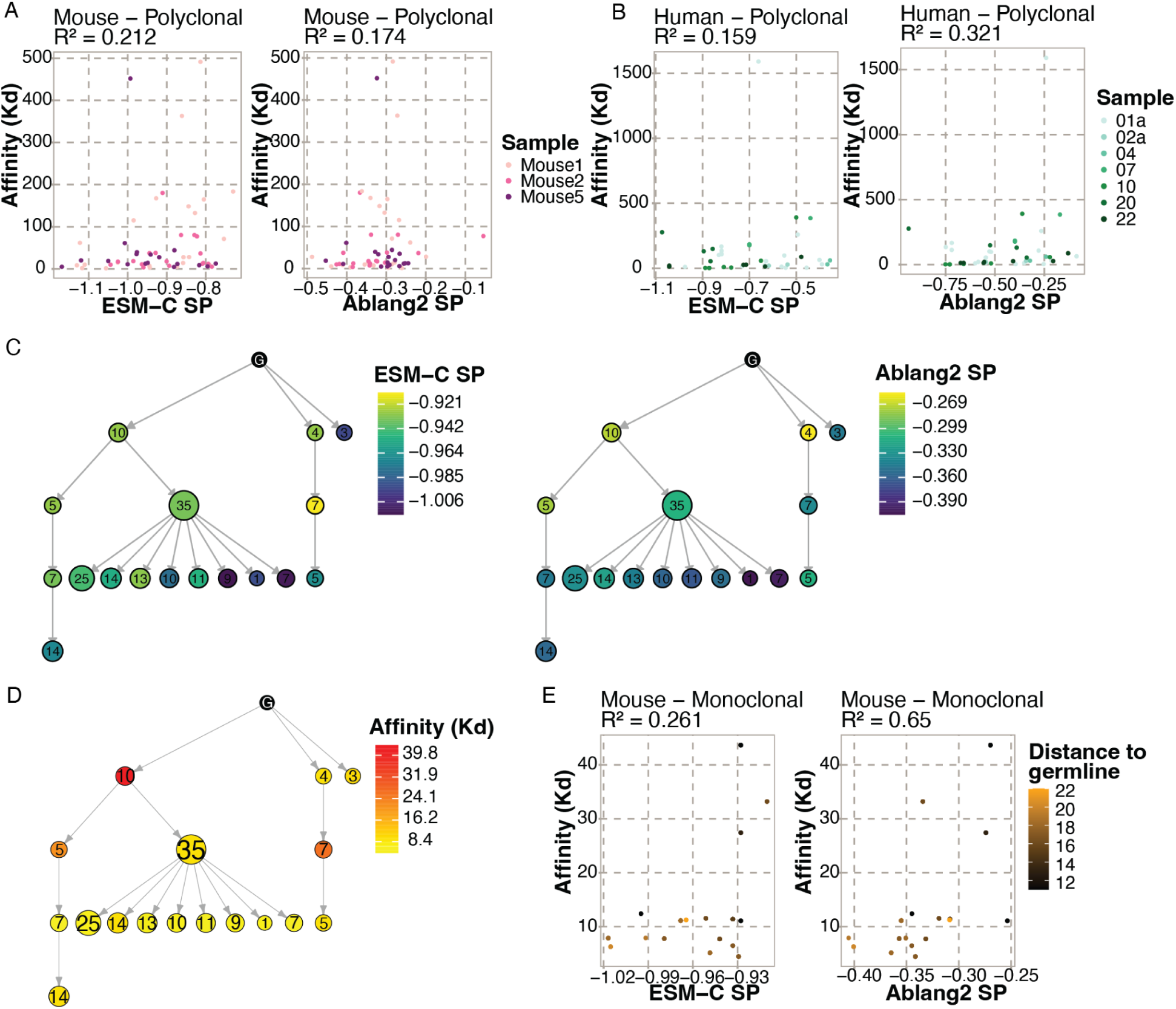
SP correlation with binding affinity. A) Spearman Correlation between polyclonal binding affinity against Ovalbumin and heavy chain ESM-C SP and Ablang2 paired chain SP for three mouse samples. B) Spearman Correlation between polyclonal binding affinity against SARS-Cov-2 S protein and heavy chain ESM-C SP and Ablang2 paired chain SP for seven individuals from Kim et al. (27) C) Lineage tree of the largest IgG clone of Mouse1 colored by heavy chain ESM-C SP (left) and Ablang2 paired SP (right). D) Lineage tree of the largest IgG clone of Mouse1 colored by binding affinity to Ovalbumin. E) Spearman Correlation between monoclonal binding affinity to Ovalbumin and heavy chain ESM-C SP and Ablang2 paired SP for the largest IgG clones of Mouse1. The color indicates Levenshtein distance to the germline sequence.

In addition to their polyclonal affinity analysis, Neumeier et al. (9) investigated antigen binding on the monoclonal level by measuring the affinity (BLI) of the sequences in the largest binding IgG clone of Mouse1. Congruent to the polyclonal analysis, they observed no correlation between the expansion of intraclonal sequence variants and binding affinity. Since we concluded that distance to the germline was in direct correlation with SP for the human data, we questioned how this would relate to binding affinity. To this end, we calculated the SPs of these monoclonal sequences and constructed a lineage tree based on both the heavy and light chains (Figures 5D, 5E). Additionally, even in the case of Ablang2, where SP calculations involve paired heavy and light chains, we still detected a positive correlation between SP and Kd values (r^2^ = 0.65) (Figure 5E), thereby suggesting that antibodies with higher affinities correspond to lower SPs. Additionally, variants with the weakest binding affinity were shown to be closest to the germline sequence (Figure 5D). Taken together, these results indicate that SPs correlate with binding affinity on both the polyclonal and monoclonal level.

## Discussion

Inspired by the work on in vitro antibody engineering performed by Hie et al. (21), we aimed to explore the relationship between PLM-based likelihoods and in vivo features of B cell selection and evolution. We utilized general and antibody-specific PLMs on single-cell V(D)J sequencing data of humans and mice and discovered that SPs are correlated with various features of B cell selection, such as V-gene usage, isotype, and antigen specificity. These SPs also showed a correlation with features of B cell evolution, such as SHM and affinity maturation. Binding affinity data revealed an overall lower SP for antibodies with a higher affinity in both the human and mice datasets. This finding is in direct contrast with the results of Hie et al. where most of the PLM-suggested mutations in binding-antibodies increased the binding affinity, potentially by reducing the search space of plausible mutations (21). This discrepancy could be due to the difference between in vitro and in vivo settings, as in vivo B cell selection involves a number of complex interactions, such as T cell help, germinal center formation, antigen presentation of follicular dendritic cells, and B cell migration. Another possibility is that ESM-1b is increasing the stability of the antigen-binding residues since most of the proposed mutations occurred in the framework regions, which are generally less mutated in vivo compared to the CDR regions (31,32). In our exploitation of this discovery, we uncovered additional aspects of in vivo B cell selection and evolution influenced by PLMs. Labels on antigen specificity revealed SPs to be significantly higher for binding than for non-binding antibodies in the human COVID-19 dataset from Kim et al. (27). However, this difference in SP is minor and warrants further experimental exploration given the PLMs in this study are agnostic to antigen-specificity during training. The mouse dataset is from BALB/c mice housed under pathogen-free conditions, thus most expanded clones are expected to be OVA-binding. This is the case for the most expanded IgG clones, but highly expanded non-binding IgM and IgA clones were also detected. This bystander activation could be a result of age- or autoimmune-associated B cells which are observed in inbred mice strains in previous findings (33,34).

When engineering an antibody, solely altering the CDR3 region instead of the full V(D)J might be desirable to maintain protein stability while focusing on the most significant sequence region for antigen binding. However, our results indicate that PLMs are dependent on the long-range interactions of the full V(D)J sequence to extract biologically meaningful features of the BCR repertoire. We included the recently developed ESM-C in our work, which is trained on more data than its predecessor ESM-1b, and includes protein structure information. We observed SPs calculated with ESM-C to be generally lower than those from other PLMs, which is explained by a more even distribution of RLs per position. This could indicate a lower certainty of ESM-C regarding antibody sequence likelihoods or understanding a wider range of tolerated residues. Additionally, we found antibody-specific PLMs to have a higher correlation with features of B cell selection and evolution than general PLMs. A limitation of most the PLMs used here is the lack of cross-chain representation. The pairing of the heavy- and light chain is crucial for antigen binding, which shapes B cell evolution. It was recently demonstrated that training or fine-tuning a PLM on natively paired chains improved the models’ performance on antigen specificity classification tasks (35) and binding affinity prediction (36). Another consideration when using these PLMs is selection bias in the training data. In these databases, most antibody sequences closely resemble germline genes (20), which helps explain the inverse correlation between SP and distance to the germline. In addition to the germline bias in training data, we discovered a bias towards overrepresented isotypes and V-genes in the OAS. It should be noted, however, that the general PLMs in this work are not directly trained on sequences from the OAS and apply a strict sequence similarity threshold to their training data. To benefit from the paired heavy and light chain sequences in our datasets and potentially overcome the germline bias, we used the recently developed model Ablang2 in our analysis. Despite its ability to embed the heavy and light chains together and its optimization on predicting non-germline residues, we did not observe major differences between Ablang1 and Ablang2 regarding their correlation with features of B cell selection and evolution. Pre-training antibody-specific PLMs with SHM as an objective function could improve the learning of B cell evolution. Wang et al. (37) showed that incorporating antibody-specific evolution mechanisms such as germline prediction and mutation position prediction benefits antibody representation learning. Lastly, including the antigen and structural information in the model architecture might further improve the PLMs, as recently demonstrated by Shanker et al. (25).

Based on our lineage trees, we observe that PLMs routinely rank SHM as one of the most likely mutations, and mutating residues generally have a slightly lower likelihood than conserved residues. However, the inherent undersampling of single-cell immune repertoire sequencing experiments makes it challenging to conclusively label conserved and mutating residues. Furthermore, epistatic mutations further complicate our results, as the presence of multiple mutations influences the likelihood of the other residues. Edges covering multiple mutations could be due to under-sampling since typically one base pair mutates per round in the germinal center (38). Computational models of SHM prediction are usually based on k-mer subsequences from 3- to 7-mers (32,39), and such studies have suggested that a longer k-mer better captures SHM as it considers a wider neighborhood (40–42). For this reason, PLM-based RLs could be a valuable option for SHM prediction, as this considers the context of the entire input sequence. The potential to incorporate factors such as activation-induced cytidine deaminase and polymerase hotspots, structural features, and G-content during fine-tuning of foundational pre-trained models using low-rank adaptation could improve SHM prediction accuracy (40).

Together, our results emphasize the ability of PLMs to represent the unique features of in vivo B cell selection and evolution. Importantly, our results demonstrate how various features, such as the model, training data, and contextual information, can dramatically influence PLM output likelihoods. While the promise of leveraging PLMs for in vitro antibody engineering has been demonstrated, further work is needed to fully leverage PLMs to predict functional features of antibody selection in vivo.

## Methods

Code to reproduce all results in this manuscript can be found in the following GitHub repository: https://github.com/dvginneken/PLM_Likelihoods

### Data Collection

For this manuscript, four publicly available datasets were used. The mouse dataset was downloaded from Neumeier et al. (9) and processed using cellranger v7. The dataset corresponding to Individual1, as described in Horns et al. (7), was also downloaded from the Platypus Database with project ID “horns2020a”. Since this dataset consists of multiple technical replicates, only the sample “Influenza.vac.11.12.human.S1” was used here. The dataset corresponding to Individual2, as described in Bruhn et al. (26), was downloaded from the NCBI Sequence Read Archive using SRA Toolkit with ID SRR24140776. This data was then processed with the CellRanger VDJ pipeline v8.0.0 from 10x Genomics. The dataset corresponding to Individual3-5, as described in Kim et al. (27), was also downloaded from SRA and processed with the CellRanger VDJ pipeline. SRR IDs used were: SRR17729703 for Individual3; SRR17729692 for Individual4; SRR17729726 for Individual5. For validation of our results, we used the human data of SRR17729673, SRR17729674, SRR17729681, SRR17729714, and SRR17729725 from the Kim et al. dataset (27) and the mice data of Mathew et al. (11).

### Repertoire Analysis

CellRanger output from all datasets was further processed with the Platypus package (version 3.6.0) in R to obtain BCR repertoire information such as clonotypes, expansion, germline sequences, isotype, and v/d/j-gene usage (28). We applied the following parameters to the VDJ_build() function: trim.germlines = T, remove.divergent.cells = T, complete.cells.only = T. These parameters trim the germline to the V(D)J region, remove cells with more than 1 V(D)J transcript, and remove cells missing either a heavy or light chain transcript. Next, cells without an annotated germline, heavy chain C-gene, or duplicated barcode were removed for further analysis. Somatic hypermutations were quantified by aligning each sequence to its corresponding germline and calculating the Hamming distance.

### Protein Language Models

The following six pre-trained PLMs were used in this manuscript: ESM-C (43), ESM-1b (12), ProtBERT (13), Ablang1 (18), Ablang2 (20), and Sapiens (22) using the Transformers package (version 4.36.1) in Python. All the heavy and light chain amino acid V(D)J and CDR3 sequences were used as input. For Ablang1 (18) and Sapiens (22), the separate heavy and light chain model was used, corresponding to the chain of the input sequence. As output from these models, we got the per-sequence pseudolikelihood and a probability matrix of the per-residue likelihood. The per-sequence pseudolikelihood (SP) was calculated by summing up the log-scaled probability of every residue and dividing this by the sequence length. For input source CDR3-from-VDJ, the entire V(D)J sequence was used as input for the PLM, but only the CDR3 sequence was used to calculate the pseudolikelihood.

### Constructing Lineage Trees

Phylogenetic networks of the clonal lineages were created of the heavy chain sequences using the default algorithm and maximum parsimony algorithm of the R-package AntibodyForests (v1.0.0) (44).

### Analysis of SHM in Lineage Trees

Substitutions along the edges of the lineage trees were identified by retrieving the unique sequences of all nodes of all trees except the germlines. If the sequence before the edge (seq1) and the sequence after the edge (seq2) had the same length, differential positions were identified. Next, a PLM probability matrix was computed for seq1 (the probability of each amino acid per position). For each differential position, the probabilities were ranked in descending order and the rank of the amino acid at this position in seq2 was selected as the substitution rank. If there were multiple differential positions for an edge, the average was computed as the substitution rank. For the predictability of the SHM location, we did the same workflow. However, we selected the rank of the amino acid in seq1 instead of seq2. In addition to the rank, we also analyzed the average probability (likelihood) of the differential- and identical positions of seq1.

### OAS Statistics

To obtain statistics on the sequence composition of OAS, we downloaded files containing human and mouse sequences separately on 29.04.2025, and extracted isotype labels and the number of unique sequences from the file headers. For all human heavy-chain sequences, we then counted the number of sequences per V-gene family based on the “v_call” column.

## Acknowledgments

We thank Victor Greiff for his insightful feedback on the manuscript, which greatly improved its clarity and quality.

## Supplementals

**Figure S1.**
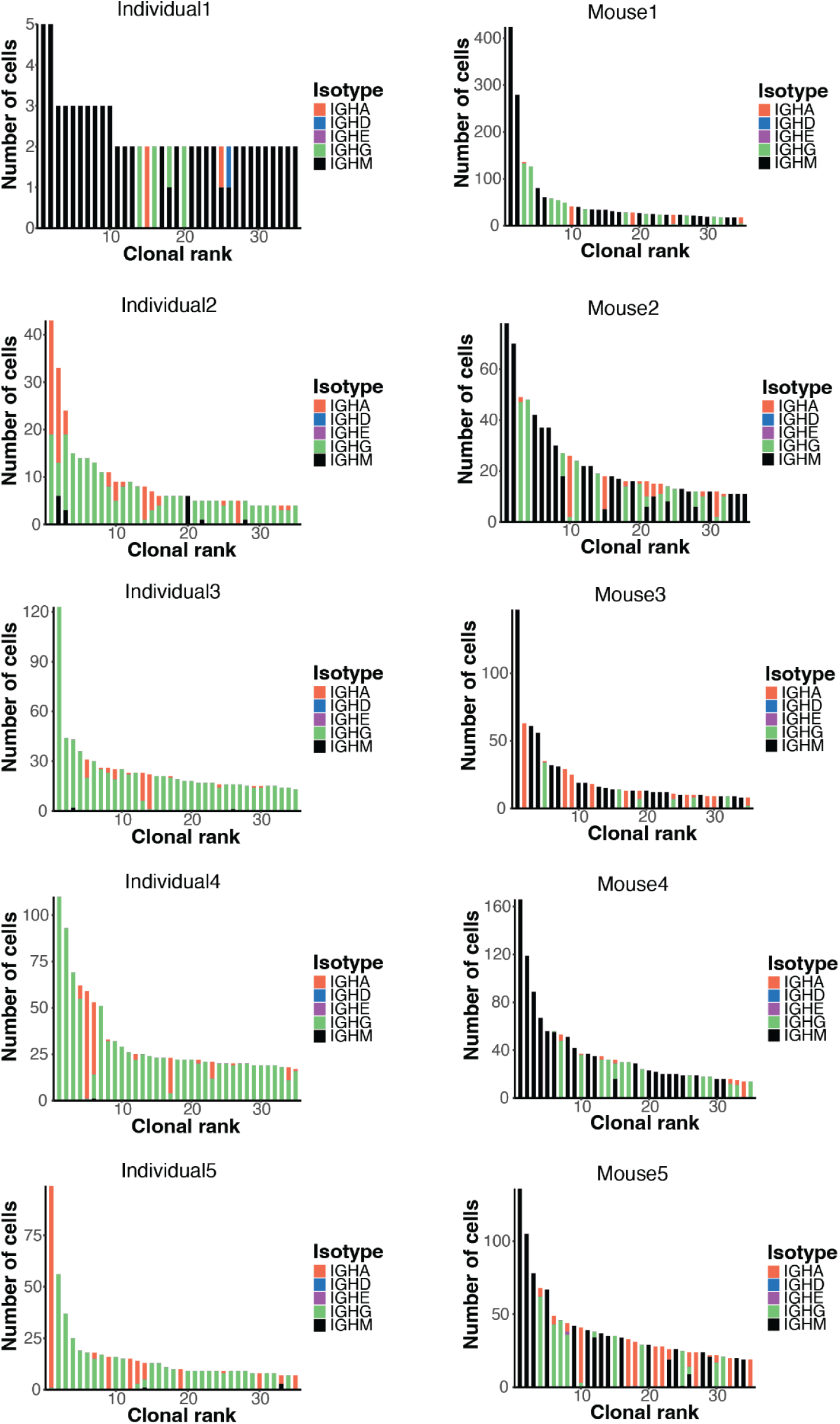
Clonal expansion of all human samples (left) and mouse samples (right) colored on isotype. Ranked bar plots depict the thirty-five most expanded clones of each sample.

**Figure S2.**
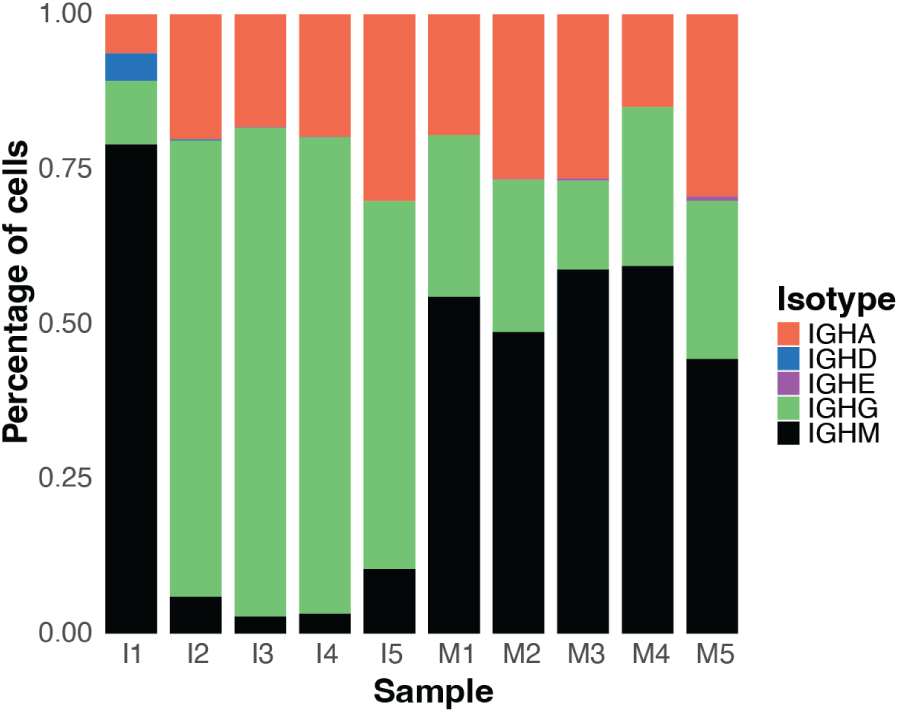
Isotype distribution per sample. I = Individual, M = Mouse.

**Figure S3.**
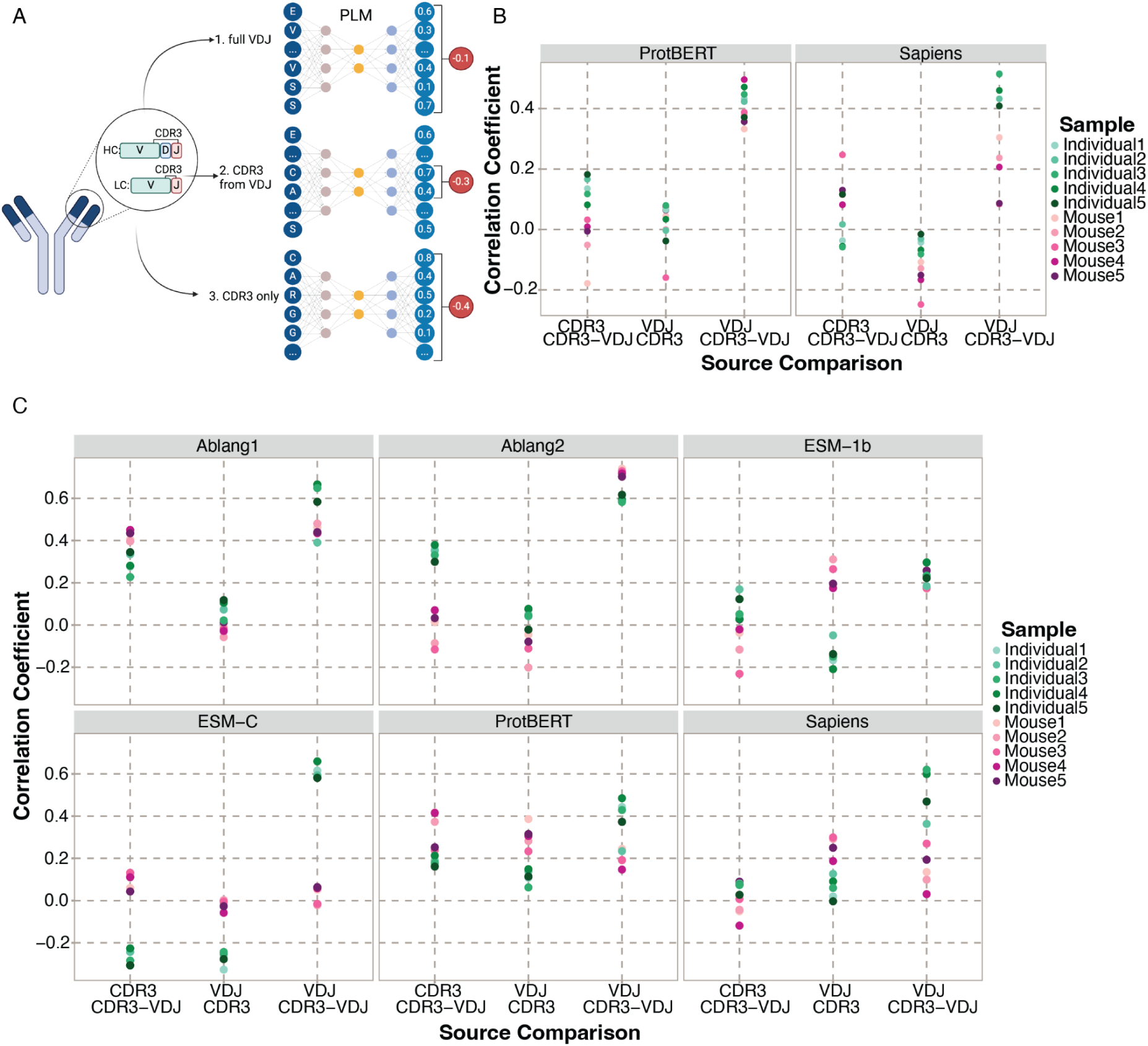
Input source correlations. A) Schematic overview of the three input sources; 1: full VDJ as input and full VDJ used to calculate SP, 2: full VDJ as input but only CDR3 used to calculate SP, 3: CDR3 as input and CDR3 used to calculate SP. Created in BioRender. B) The Pearson Correlation of SPs of the heavy chains between the three sources of input for ProtBERT and Sapiens (other four PLMs are in main Figure 1C). C) The Pearson Correlation of SPs of the light chains between the three sources of input for all PLMs..

**Figure S4.**
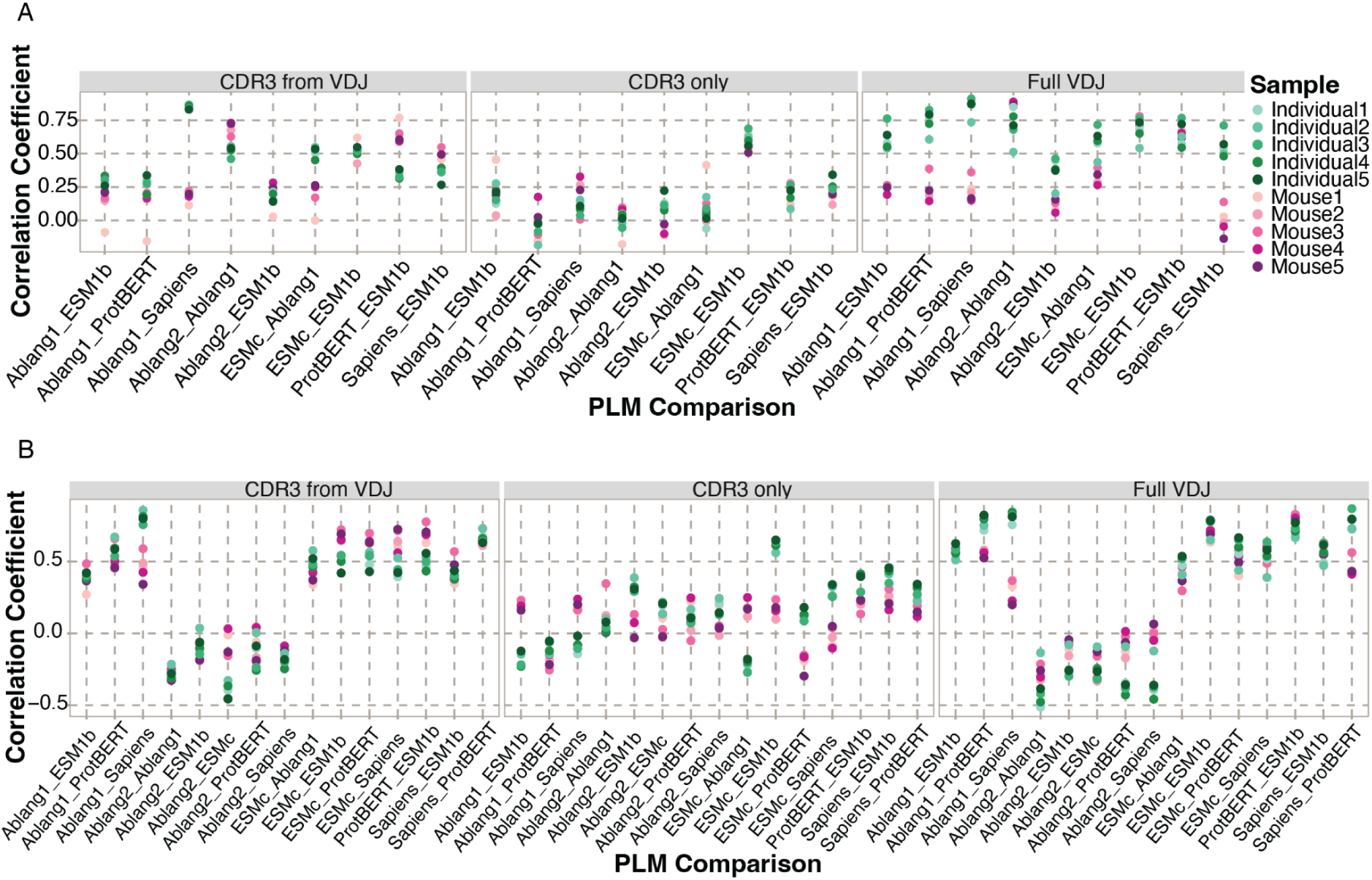
Pearson Correlation between SPs of different PLMs. A) Correlation between SPs of the heavy chains calculated with different PLMs for each of the three input sources. B) Correlation between SPs of the light chains calculated with different PLMs for each of the three input sources.

**Figure S5.**
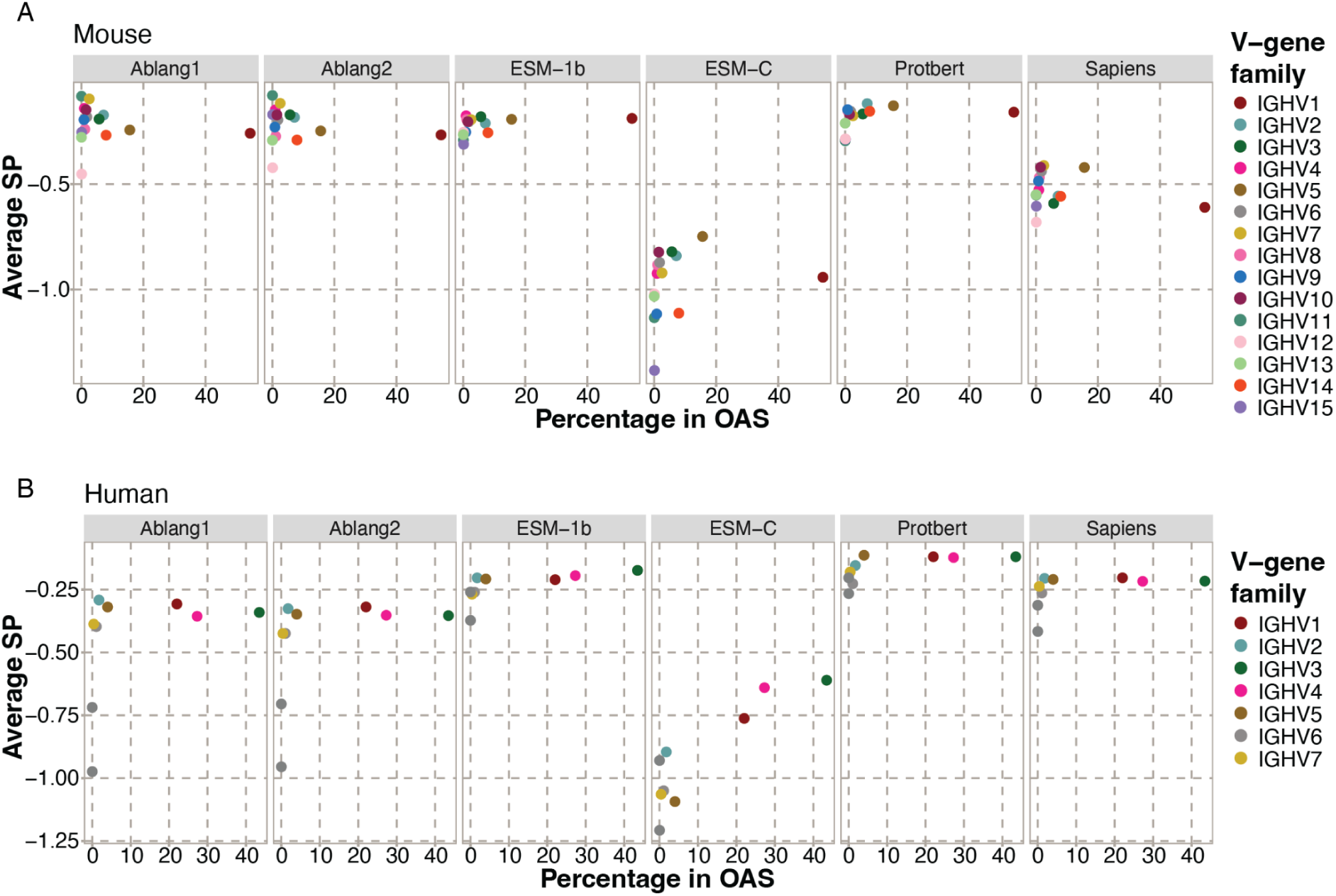
Correlation between SP and isotype frequency in the OAS. A) The average SP per V-gene family and the percentage of unique sequences of this V-gene family in the OAS database for the mouse samples. B) The average SP per V-gene family and the percentage of unique sequences of this V-gene family in the OAS database for the human samples.

**Figure S6.**
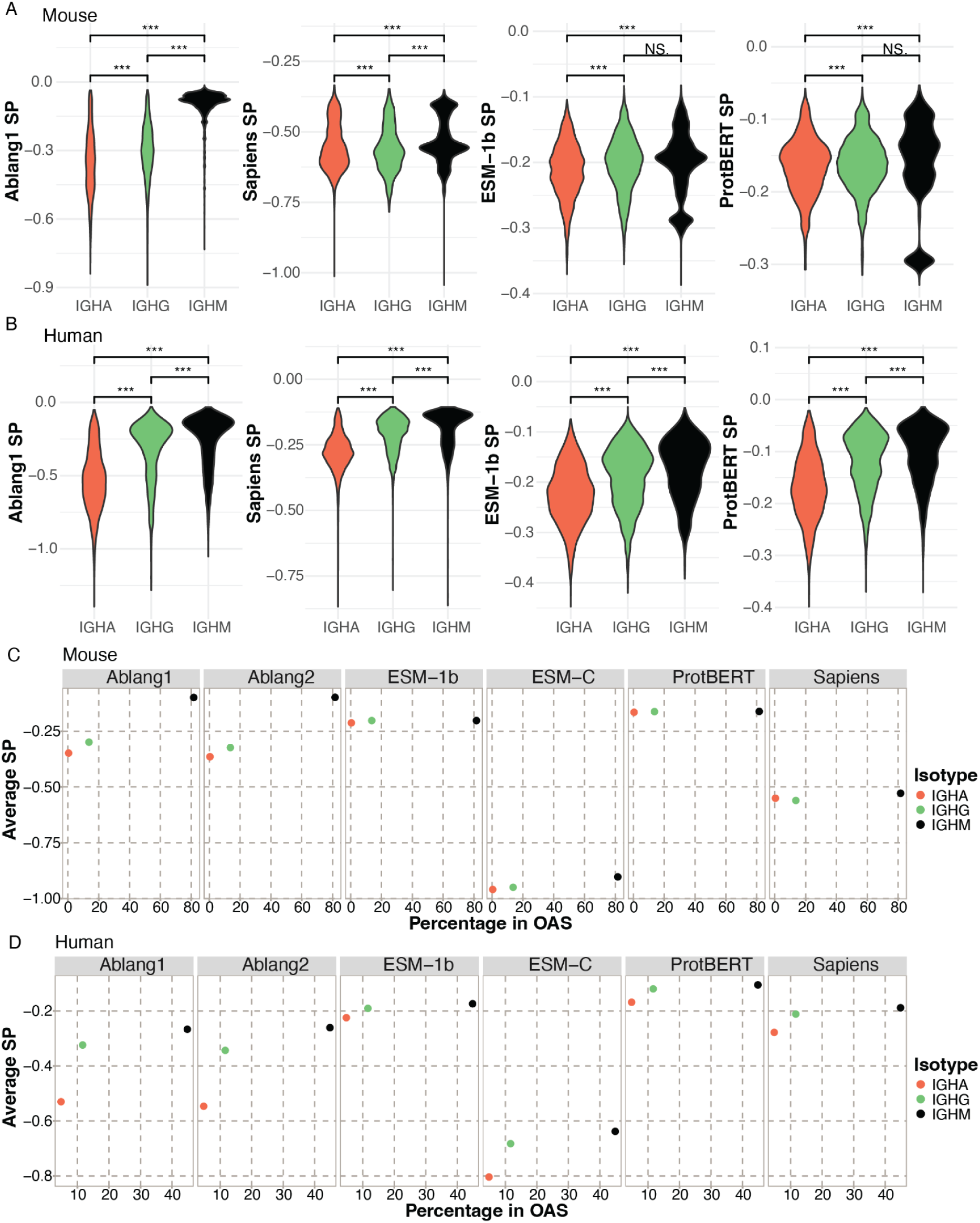
Correlation between SPs and isotypes. A) Distribution of SPs of mouse BCRs from certain isotypes. B) Distribution of SPs of human BCRs from certain isotypes. T-test significance: *** = adjusted p-value below 0.001, NS = non-significant. C) The average SP per isotype and the percentage of unique sequences of this isotype in the OAS database for the mouse samples. D) The average SP per isotype and the percentage of unique sequences of this isotype in the OAS database for the human samples.

**Figure S7.**
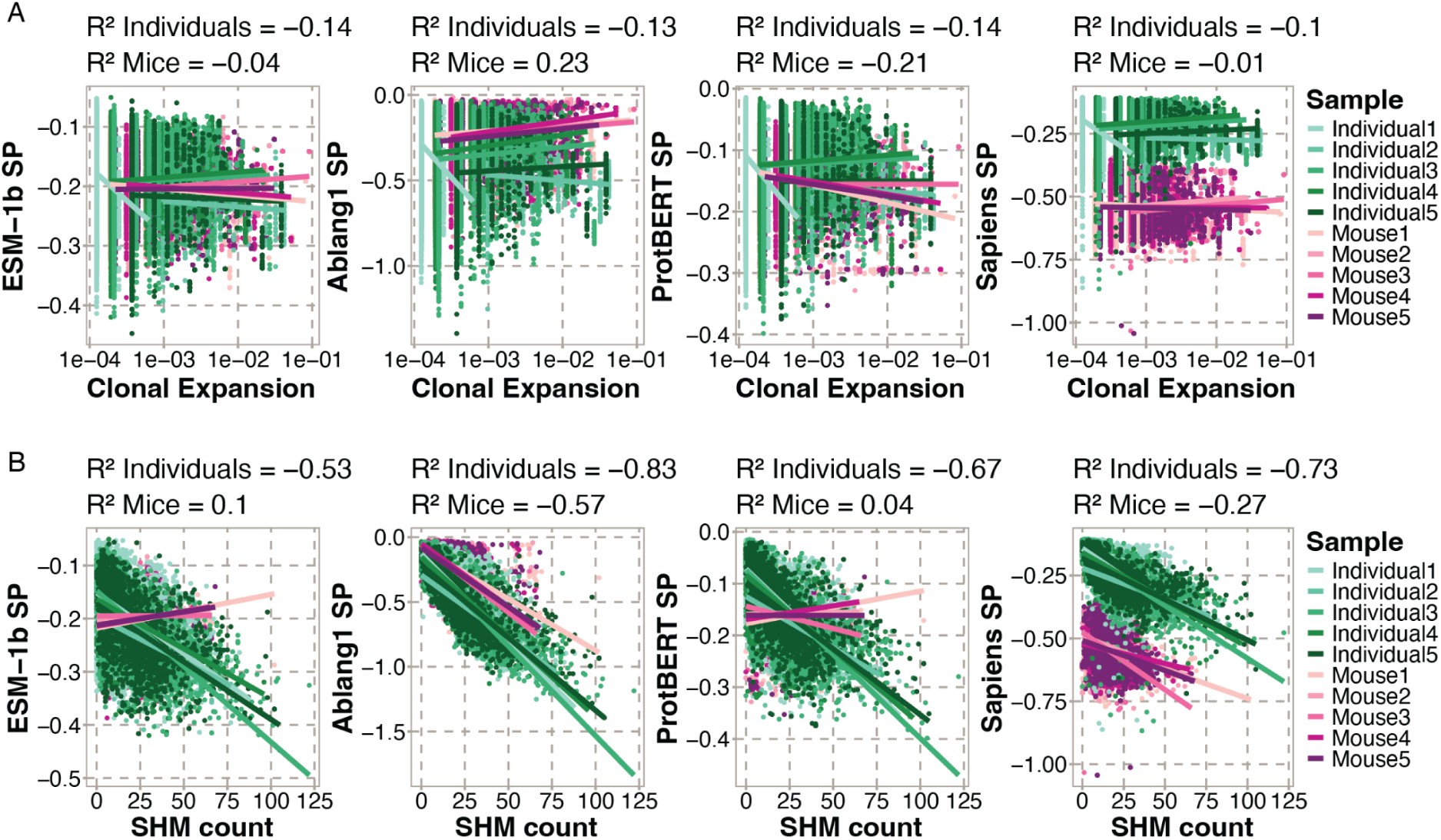
Correlation between SPs and features of B cell repertoire evolution. A) Pearson Correlation between normalized clonal expansion (number of cells per clonotype divided by the sample size) and SP for four PLMs. B) Pearson Correlation between the amount of SHM (hamming distance from the germline) and SP for four PLMs.

**Figure S8.**
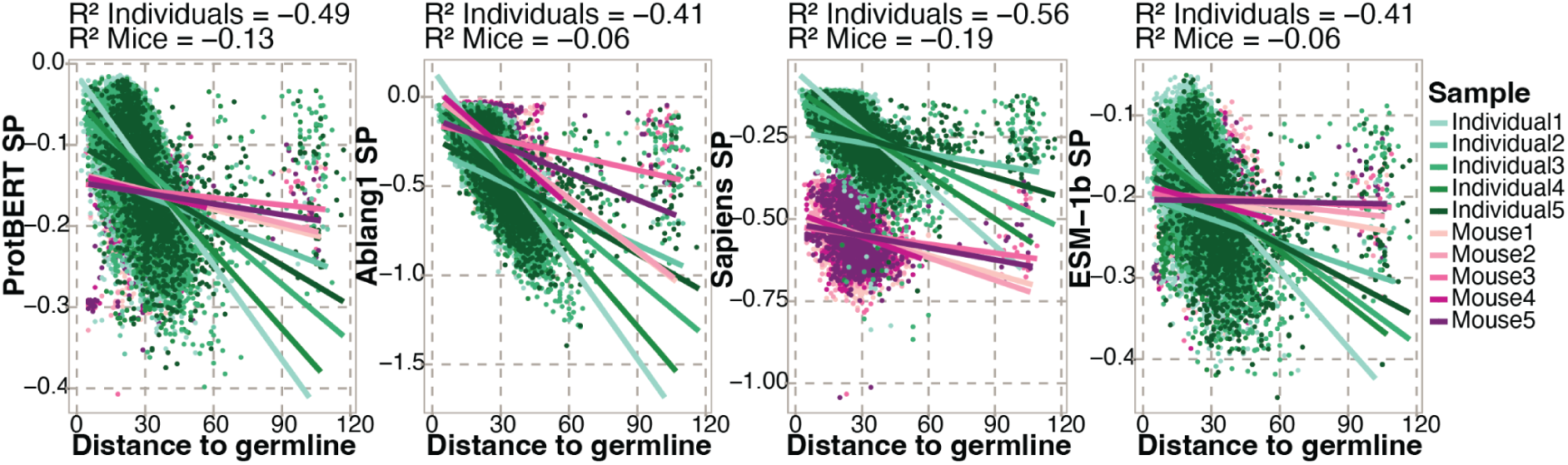
Correlation between SP and the sum of edge lengths to the germline for each heavy chain BCR sequence in the lineage trees of all samples.

**Figure S9.**
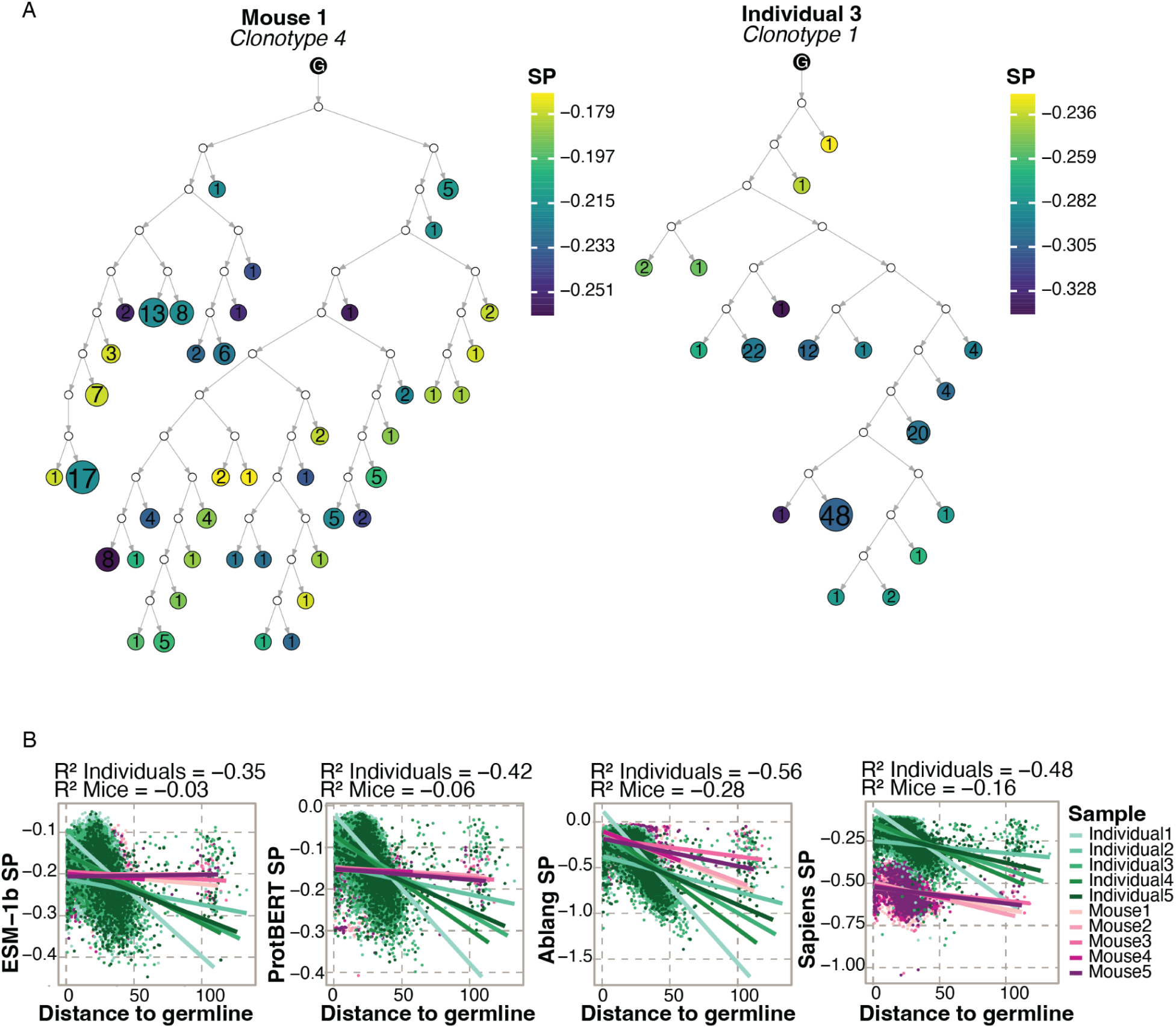
Phylogenetic analysis of the clonal lineages constructed with a maximum parsimony algorithm. A) Representative plots of the lineage trees colored by ESM-1b full-VDJ SP. White nodes represent intermediate nodes recovered by the phylogenetic algorithm. B) Correlation between SP of all PLMs and the total edge length (Levenshtein distance) to the germline for each sequence in all trees.

**Figure S10.**
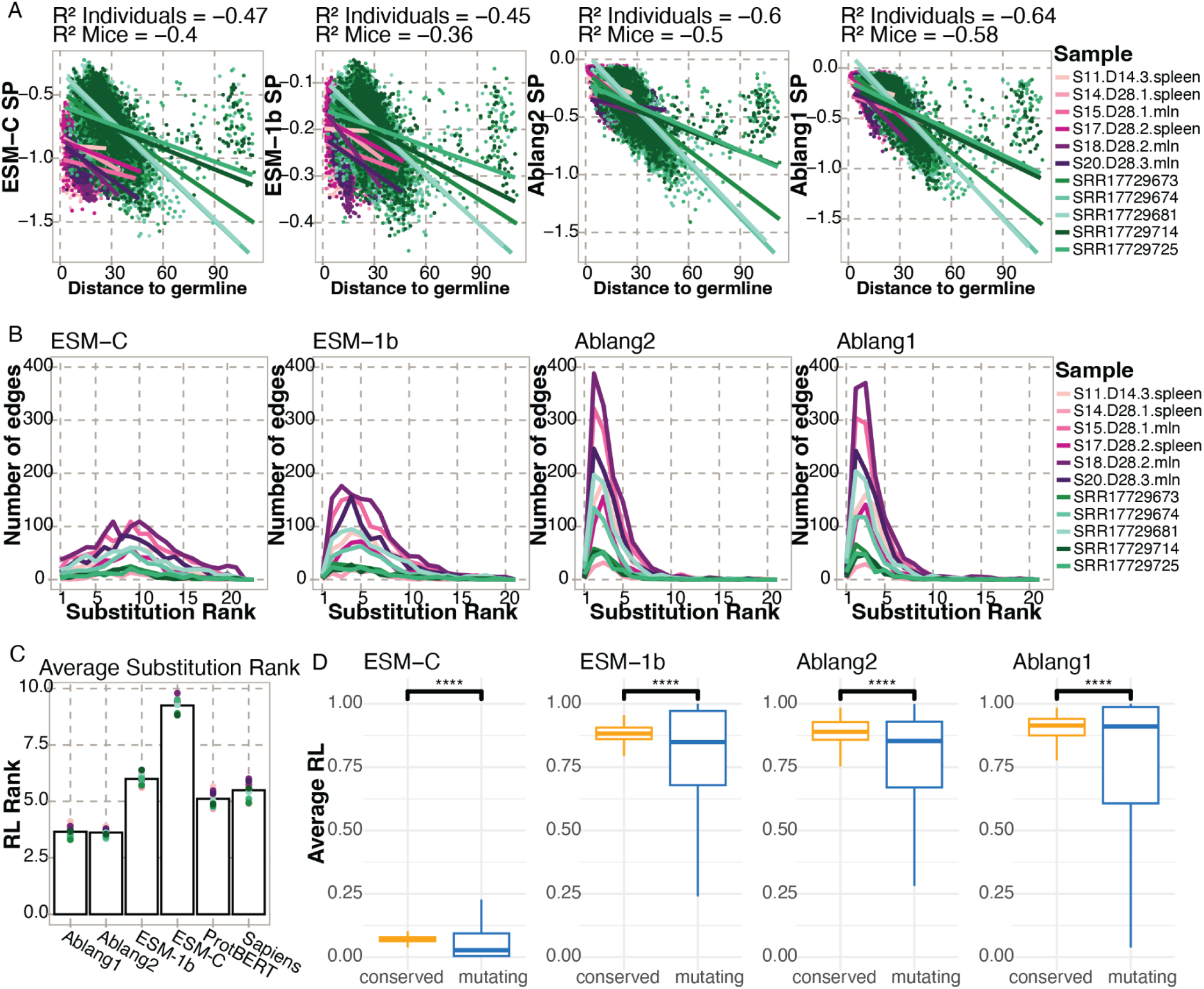
Additional data for lineage tree validation. Purple indicates mouse samples from Mathew et al. Cell Rep, 2022. Green indicates additional human samples from Kim et al. Nature 2022. A) Pearson Correlation between SP and the total edge length to the germline (Levenshtein distance) for each sequence in all trees. B) The RL ranks of the substitutions along the edges of the lineage trees. The average rank is used for edges with multiple substitutions. C) Mean substitution RL rank for each sample (dots) and average of all samples (bars) per PLM. E) Difference in average RL between conserved and mutating residues. T-test significance: **** = adjusted p-value below 0.0001.

**Figure S11.**
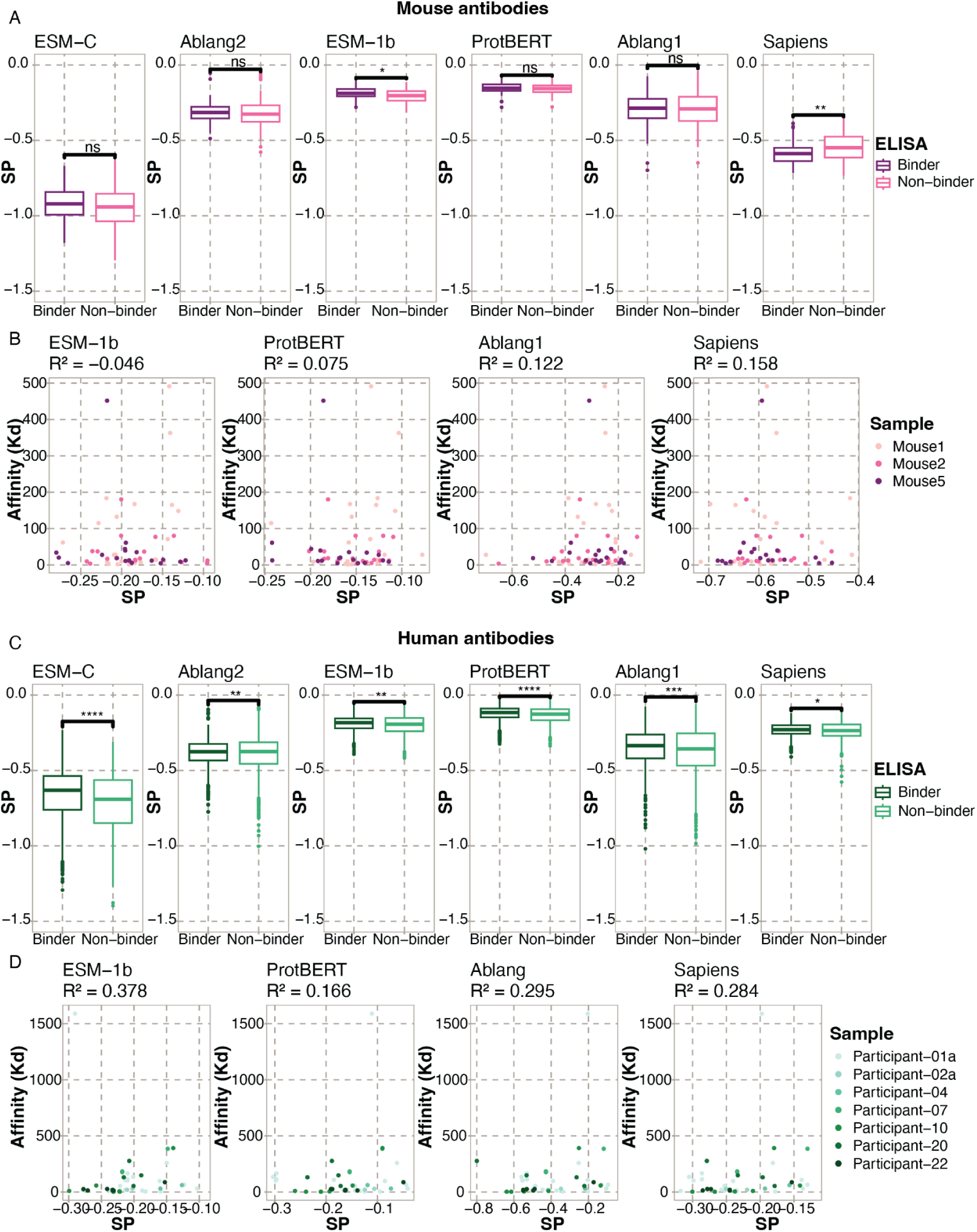
Spearman Correlation of SP with polyclonal antigen-specificity and binding affinity. A) SP and antigen-specificity against Ovalbumin. B) Correlation between binding affinity and SPs for mouse samples. C) SP and antigen-specificity against Sars-COV-2 S protein. D) Correlation between binding affinity and SP for 7 individuals (sample IDs correspond to those of Kim et al.(26)). (T-test significance: ns = p > 0.05; * = p ≤ 0.05; ** = p ≤ 0.01; *** = p ≤ 0.001; **** = p ≤ 0.0001)

## Notes

https://github.com/dvginneken/PLM_Likelihoods

### Competing Interest Statement

The authors have declared no competing interest.

### Summary of Updates

The addition of newer PLM models (both general and antibody-specific). The antibody-specific PLMs additionally can calculate pseudolikelihoods for paired heavy and light chains. Additional repertoires were included to better understand the ability to generalize our findings, highlighting which residues within the BCR are likely to mutate and to which amino acid. Mining of the OAS database to understand potential biases in training data and how they relate to the performance of models.

